# Designing and studying a mutant form of the ice-binding protein from *Choristoneura fumiferana*

**DOI:** 10.1101/2020.08.31.275651

**Authors:** Ksenia A. Glukhova, Julia D. Okulova, Bogdan S. Melnik

**Author notes:** **Corresponding Author:** Bogdan S. Melnik.

## Abstract

Ice-binding proteins are expressed in the cells of some organisms, helping them to survive extremely low temperatures. One of the problems in study of such proteins is the difficulty of isolation and purification. For example, eight cysteine residues in cfAFP from *Choristoneura fumiferana* (the eastern spruce budworm) form intermolecular bridges during the overexpression of this protein. This impedes the process of the protein purification dramatically.

In this work we designed a mutant form of ice-binding protein cfAFP, which is much more easy to isolate that the wild-type protein. The mutant form named mIBP83 did not lose the ability to bind to ice surface. Besides, observation of the processes of freezing and melting of ice in presence of mIBP83 showed that this protein affects the process of ice melting, increasing its melting temperature, and at least does not decrease the freezing temperature.

## Introduction

Spatial structure of ice-binding protein cfAFP from *Choristoneura fumiferana* was obtained in 2000 by nuclear magnetic resonance method [1]. Later, its structure was refined by X-ray analysis in 2002 [2]. The structural experiments elucidated that the protein comprises a triangular prism formed by beta-strands (Figure 1). Threonine residues on one of the faces of this prism form an ice-binding surface (Figure 1 a, b) [1–3].

**Figure 1.**
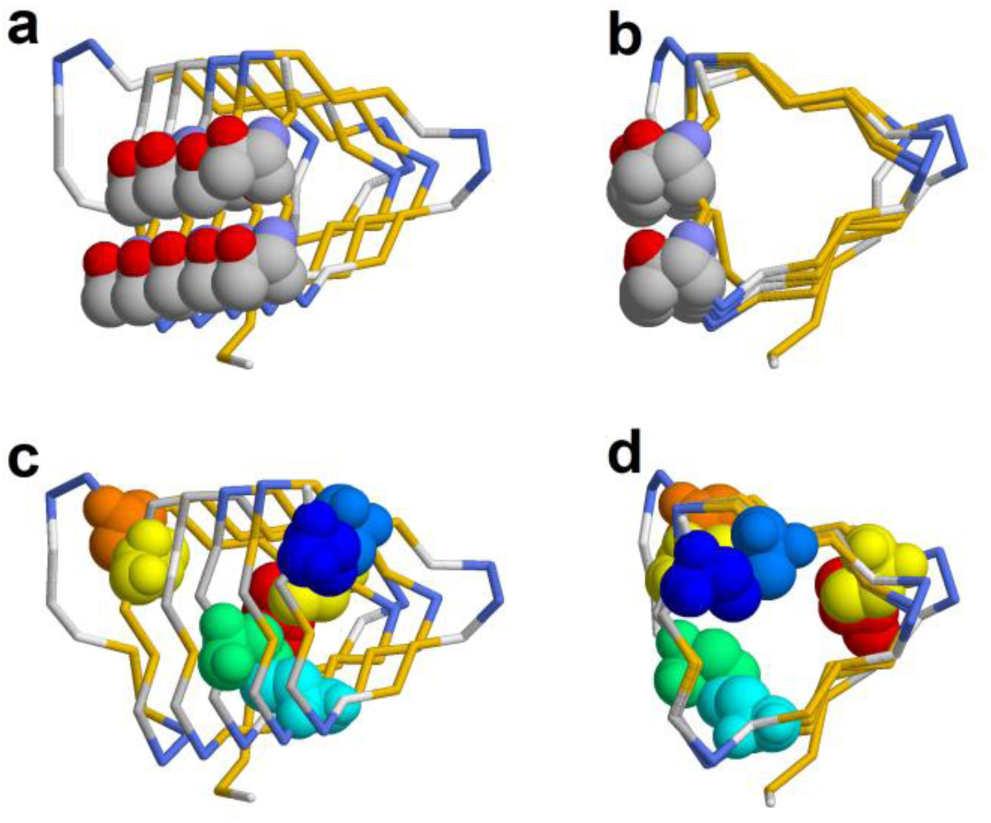
Ice-binding protein cfAFP (PDB ID: 1L0S). The polypeptide chain is displayed as a wire model. Ice-binding threonine residues are shown as 3D spheres on projections (a) and (b). Cysteine residues are shown as 3D spheres on projections (c) and (d). Eight cysteine residues form four disulfide bonds.

Besides, eight cysteine residues forming four disulfide bonds are present in the amino acid sequence of cfAFP (Fig. 1 c, d).

The ability of cfAFP protein to bind to ice surface was studied by several research groups [3,4]. Also, the ability of this protein to change the melting point of ice and freezing point of solution (hysteretic properties of the protein) was studied [5–7].

On the earlier stages of our studies, we met the challenges during isolation and purification of the protein. Beside the overexpression of the target protein in *E. coli* cells, only a small amount of the protein can be isolated and purified (about 0.5mg per liter of cultural medium). The protein formed intermolecular disulfide bonds at different steps of isolation and purification, and due to that it fell into inclusion bodies or formed a pellet. Addition of the reducing agents (mercaptoethanol or DTT) helped at the early stages of purification, but the intermolecular bonds were still formed when the concentration of reducing agents was decreased at the final stages of purification.

## Results

### Design of the mutant protein

After analysis of cfAFP protein (Figure 1), we decided to design its mutant variant which would be less susceptible to aggregation during isolation, but would retain the ability to bind to ice surface, specific for the wild-type protein. For this purpose, we substituted six of eight cysteine residues and truncated the protein. We supposed that one disulfide bridge would be enough for protein structure stabilization, whereas substitution of six cysteines should decrease the probability of intermolecular bond formation during isolation and purification. Since cfAFP protein consists of repeated turns (Figure 1), the amino acid substitution can be simply chosen based on the amino acid composition of the adjacent protein turns. Moreover, we designed the substitutions increasing the regularity of the protein structure. A total of nine substitutions were performed: C4V, C17V, C67A, C80V, I68T, T66G, T49K, C62V, C85V (numbering according to PDB ID: 1L0S). Figure 2 demonstrates the protein structure with all the amino acid substitutions shown, and the C-terminal region that is deleted in case of the mIBP83 mutant or substituted for a linker in fusion protein is highlighted (described below).

**Figure 2.**
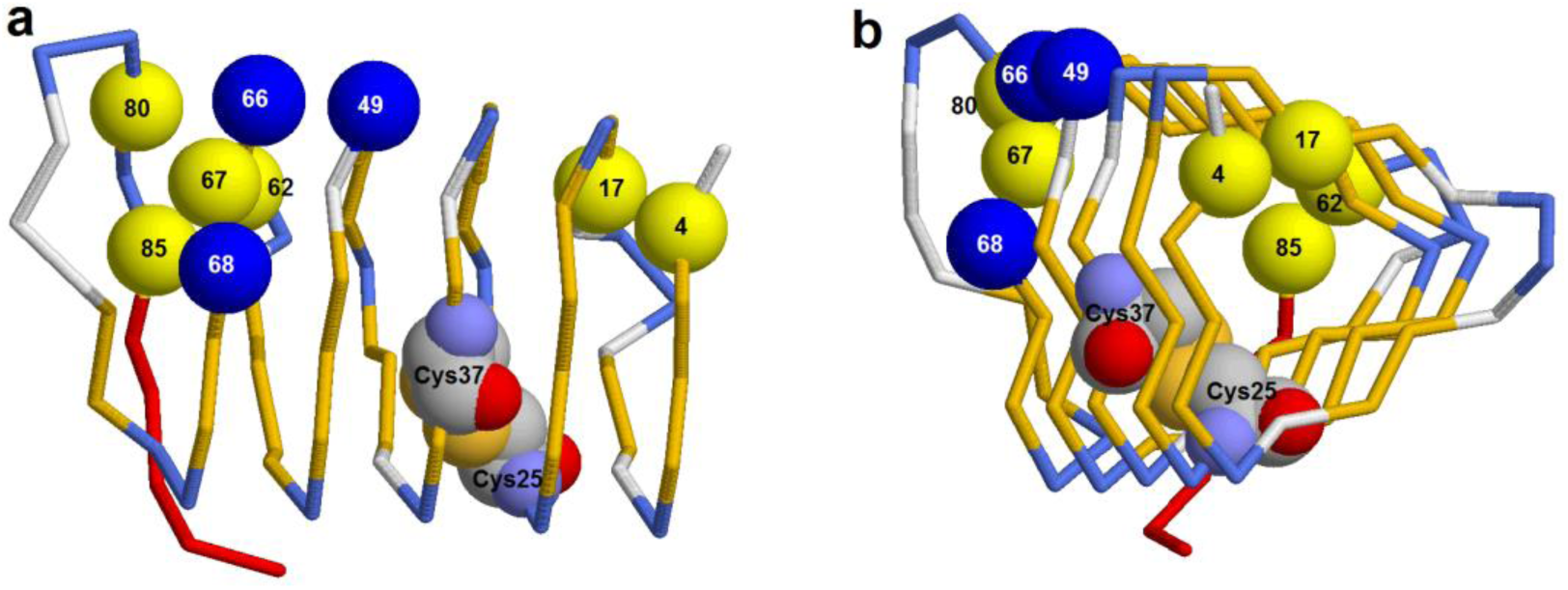
Structure of ice-binding protein cfAFP (PDB ID: 1L0S) with amino acid substitutions. Balls show Cα atoms of substituted cysteines (yellow) and other (blue) amino acids. **a** - side view; **b** — view from the N-terminus of the protein. Two cysteine residues forming a unique disulfide bridge in the mutant protein mIBP83 (Cys37, Cys25) are shown as 3D space-filling models. C-terminal region that is truncated in mIBP83 and substituted to a linker in chimeric protein mIBP83-GFP is highlighted in red.

The nucleotide sequence enconding the designed mutant protein mIBP83 was inserted into pET-22b plasmid using conventional modern methods of genetic engineering (See Materials and Methods). Besides, a chimeric protein named mIBP83-GFP was designed, where mIBP83 was fused with green fluorescent protein [8, 9] by eight-residue linker (GASGAGMA) (Cycle3-GFP protein was used, see Materials and Methods section). Such a chimeric protein is required to visualize the binding of mIBP83 to ice surface.

The designed mutations dramatically decreased the number of possible intermolecular cysteine bridges. Mutant proteins mIBP83 and mIBP83-GFP were easily isolated by conventional methods in large amounts without addition of reducing agents (See the details of isolation procedure in Materials and Methods section).

### Examination of the ability of mIBP83-GFP protein to bind to ice surface

To test the ice-binding ability of mIBP83-GFP fusion protein, the following experiment was carried out. Two identical tubes were filled with buffer solution (1.0ml, 20 mM sodium phosphate buffer, pH = 7.0) and frozen at −20°C, then incubated at room temperature till the beginning of ice melting. Thus, each vial had a piece of ice surrounded by a small volume of water. Then, mIBP83-GFP solution was added to one tube, and GFP solution was added to another one (200 µl, 2mg/ml). The tubes were irradiated by a transilluminator. If mIBP83 as a part of the chimeric protein would not lose its ability to bind to ice surface, it should cover the piece of ice floating in the tube, and the contour of this piece should fluoresce more intensely than the solution. GFP lacks the ability to bind to ice, so it would not be seen on the contour of the ice piece. Figure 3 shows the photographs of one of such experiments.

**Figure 3.**
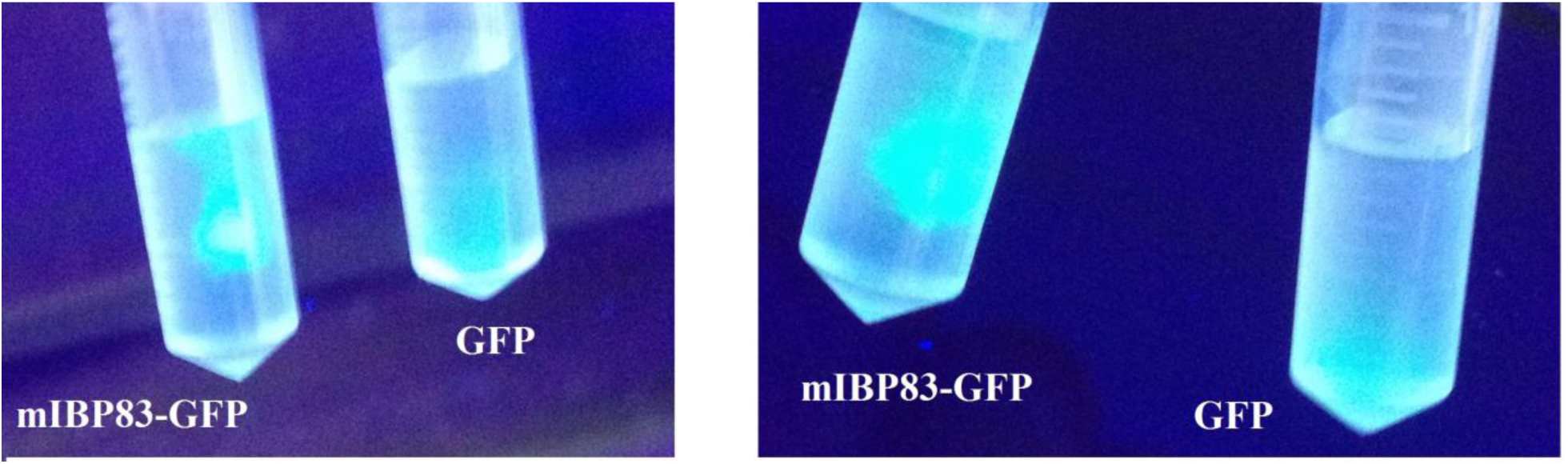
Binding of mIBP83-GFP to the ice surface. A tube with mIBP83-GFP solution is on the left, a tube with GFP solution is on the right on each image. The tubes were irradiated with ultraviolet light from below.

It can be seen in Figure 3 that the main fluorescing component in the tubes with mIBP83-GFP is the piece of ice, its contours can be distinguished. In tubes with GFP, the solution luminesces mainly in the bottom, because the tubes are irradiated from below.

Thus, one can conclude that the protein mIBP83 attached to GFP by a linker did not lose the ability to bind to ice surface.

### Examination of the ability of mIBP83 protein to affect freezing and melting points of ice

To test the ability of mIBP83 to change the freezing point of water, the following experiment was carried out. 50, 100, 150 or 200µl of solution were placed into 200 µl tube, a probe of electronic thermometer was immersed into the middle of the test tube, and it was closed. Then the tube was placed into a metal holder that was cooled by circulating liquid of the thermostat. The scheme of the instrument is shown in Figure 6 in Materials and Methods section. The thermostat cooled the tube changing the temperature linearly with 0.24 degree per minute cooling rate. If the processes of heat consumption and generation do not occur in the sample, then the temperature of sample should change linearly with time at about the same rate as in the whole thermostat. During the freezing, crystallization process occurs, heat is evolved and the temperature of the sample becomes higher. Such a peak of temperature during water freezing can be detected by a thermometer.

Figure 4 shows the curves of temperature changes in the samples depending on cooling time. The curves are fit by freezing time (shown by an arrow, Fig. 4)

**Figure 4.**
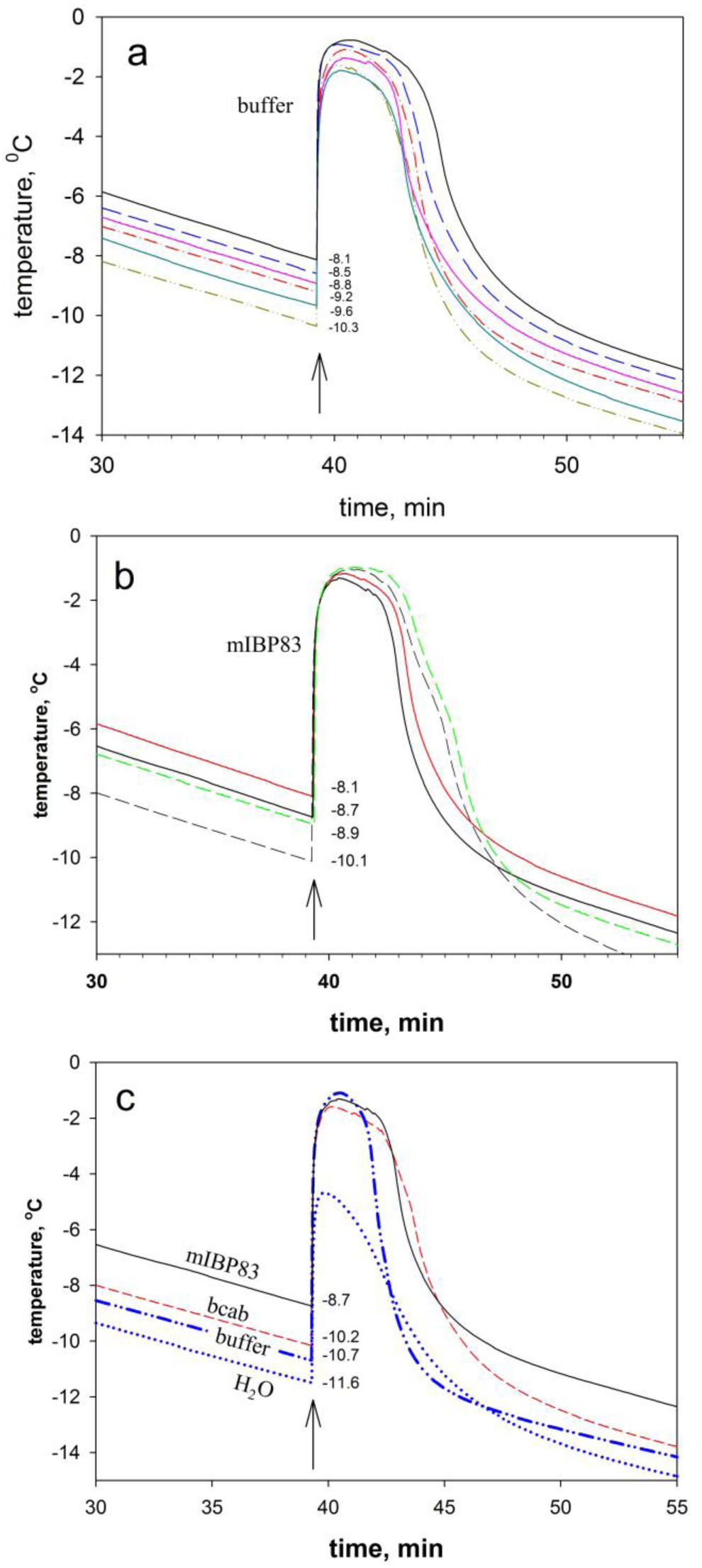
Dependence of sample temperature on cooling time. a **—** an example of six experiments of freezing 20 mM sodium phosphate solution (pH = 7, V = 150 ul); **b**-an example of four independent experiments of freezing mIBP83 solution (0.6mg/ml, 20 mM sodium phosphate, pH = 7). Solid lines — sample volume 150ul, dashed lines – sample volume 200ul. **c**-curves of cooling mIBP83 solution (0.6mg/ml, 20 mM sodium phosphate, pH = 7) (solid line), carbonic anhydrase solution (0.6mg/ml, 20 mM sodium phosphate) (dashed line), sodium phosphate buffer solution (dash-dotted line) and distilled water (dotted line). The curves were aligned by solution freezing time. The moment of freezing is shown by the arrow. Temperature values of the beginning of crystallization are written on the plot.

Figure 4a displays the cooling curves of sodium phosphate buffer. Two features can be noticed: 1) The freezing points of all the samples are far below 0 °C; 2) the experiments are unstable, i.e. the variance of freezing point values is high. The experiments plotted in Figure 4a are in accordance with the well-known fact that water can exist for a long period in overcooled state. The start of its crystallization may depend on presence of particles of dust and irregularities of walls that increase the probability of formation of ice nuclei. Presence of particles of dust and microscopic wall scratches is hard to control, thus the ideal data reproducibility is unachievable in experiments on ice freezing. Nevertheless, comparison of Figures 4a and 4b allows drawing a conclusion that presence of mIBP83 in solution does only slightly affect the freezing point of water. At least, it does not decrease it. Based on numerous experiments, one can conclude that addition of any substance (protein or buffer solution components) to pure distilled water does rather increase its freezing point. This can be explained in such way: the more components added to the solution, the harder is to avoid the presence of microparticles that fall into the solution and affect ice nucleation. Figure 4 c shows comparison of four crystallization curves of four solutions: mIBP83 solution, carbonic anhydrase (a protein that cannot be an anti-freezing agent), sodium-phosphate buffer solution and distilled water. Taking into account the instability of solution freezing experiments (Figure 4a, 4b), one can conclude that freezing of all the solutions mentioned in Figure 4c occurs almost at the same temperature. At least if there is a slight effect of each substance on freezing point of water, this effect is less than the effect of dust and irregularities, on which the ice nucleation depends.

The experiments on ice melting were much more stable. Figure 5 shows the dependence of sample temperature on heating time, i.e. melting of ice with and without mIBP83, at several different heating rates. The technical details of the experiment are described in Materials and Methods section. Figure 5a shows the scheme of all the stages of temperature measurement in a sample. At low temperatures, there is heating of ice without its melting (I in Fig. 5a). When the temperature reaches melting point, the curve begins to lag behind the «ideal» one (II in Fig. 5a) because of heat consumption during ice melting. A value shown by a real thermometer immersed into the middle of a test tube in experimental appliance will always slightly lag behind the temperature of tube walls and average temperature of the whole sample. It is because of irregular and not instant heating of the sample. If the sample volume is high enough, then the curve will become flat for a certain time when reaching the melting point (III in Fig.5a). The larger the sample volume, the longer would the curve keep flat, because ice-water mixture maintains its zero temperature. When most part of ice is melted, water is heated rapidly to the temperature of the thermostat (IV in Fig.5a). The last part of the curve (V in Fig. 5a) should reflect the process of water heating with the rate equal to the thermostat heating rate.

**Figure 5.**
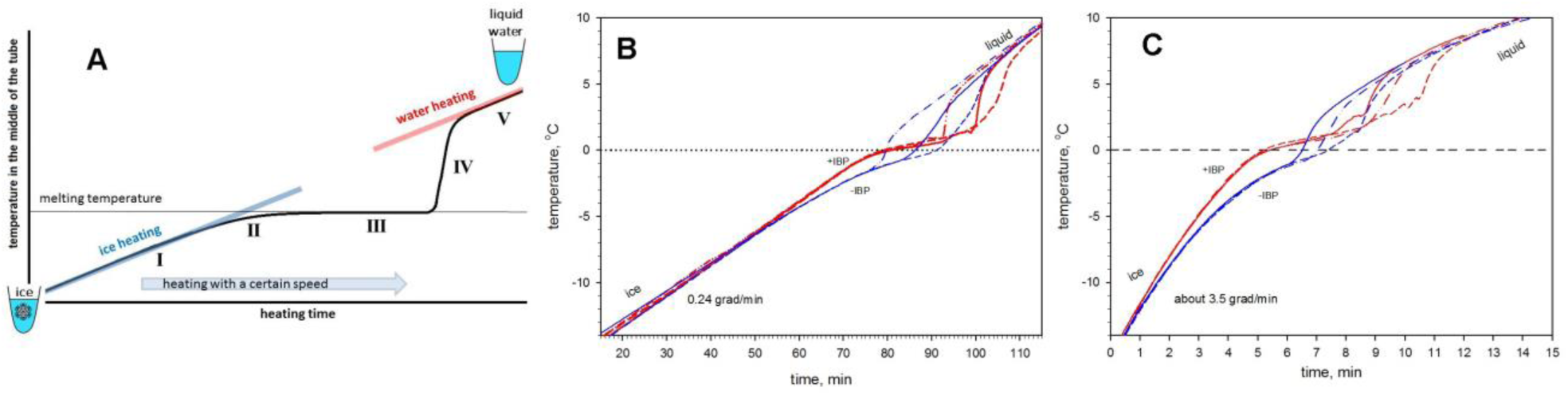
Ice melting experiments. **A** — simplified explanation of experimental curve shape: I, ice heating; II, beginning of ice melting; III, water-ice mixture maintains the melting temperature; IV, rapid heating of the liquid sample when ice is completely molten; V, water heating (see the details in the body of the manuscript). **B** and **C** - experimental curves of sample melting with different heating rate (**B** - 0.24; **C** - 3.5 degrees per minute). Red curves, 0.6mg/ml mIBP83; blue curves, samples without mIBP83. Dash-dotted line, solid line and dashed line show the curves for different sample volumes (50, 100 and 150 microliters, respectively).

Figures 5B and 5C show experimental curves of ice melting with and without ice-binding protein mIBP83. The apparent shape of the experimental curves looks similar to the schematic curve demonstrated in Figure 5A. The only difference is that stage III is very short in the experimental samples, because sample volumes are only 50, 100 and 150µl, but it can be noticed that the duration of this period increases with sample volume. It should also be noted that some thermostats are «unable to keep» the linear change of temperature at high heating rates, as shown in Figure 5C, which results in apparent curvature of the experimental curves. It is especially clear at low (around −10 °C) and high (around +10°C) temperatures. Figures 5B and 5C also show that the curves of mIBP83-free samples reach the flat line around 0°C, i.e. ice melting point, whereas the mIBP83-containing samples cross the 0°C line and reach the horizontal line in the area of positive values. In other words, ice melting temperature is increased in presence of mIBP83. On average, presence of mIBP83 in solution at 0.6mg/ml concentration increases ice melting point by one degree and it does not depend on sample heating rate. In this work, we did not have the goal to determine the exact value to which presence of mIBP83 increases the melting temperature. The main task was to understand whether presence of the ice-binding antifreeze protein (mIBP83) decreases (as it is usually believed) or, on the contrary, increases the ice melting point. The experiments displayed in Figures 5B and 5C show that presence of mIBP83 increases ice melting temperature. The elevation of ice melting point canb be explained by mIBP83 binding to ice surface. Since water molecules on the ice surface are the most mobile and unstable, their stabilization by protein binding must increase ice melting temperature. If it is exactly so, the elevation of ice melting point in presence of mIBP83 should not strongly depend on protein concentration (protein amount/ice surface ratio is always increasing during ice melting), but the melting point should strongly depend on ice binding constant of the protein.

## Conclusions

In this work, mIBP83 protein was designed; it differs from the cfAFP protein from *Choristoneura fumiferana* caterpillar by nine amino acid substitutions. The designed substitutions allow avoiding the formation of intermolecular cysteine bridges, which in turn allowed to simplify the process of isolation and purification of mIBP83 protein in large amounts.

mIBP83 protein was shown to bind to ice surface and to increase ice melting temperature by around one degree. The conducted experiments showed that presence of mIBP83 in solution does not significantly affect water crystallization temperature (at least it does not decrease it).It follows from the experiments carried out, that the ice-binding protein cannot function as a "simple anti-freezing agent” like alcohols or polyols, which decrease both melting point and freezing point of the solutions.

## Materials and methods

### Genetic constructs

To express mIBP83-GFP fusion protein in mammalian cells, mIBP83 fragment of pEX-A2-SBP-T plasmid was subcloned into pTagGFP2-N vector (Evrogen) by PCR with the primers

5’ ATTCTCGAGATGAGCGTGACCAACACCAAC 3’;
5’ ATAGGATCCCCCACGCCGCTAATTTTCAC 3’.

pEX-A2-SBP-T plasmid was purchased from Eurofins Genomics (Germany).

To produce mIBP83 protein in *E.coli* cells, DNA fragment encoding mIBP83 was subcloned into pET22b vector (Novagen) by restriction digestion of pEX-A2-SBP-T with NdeI and SalI.

To produce mIBP83-GFP fusion protein in *E. coli* cells, DNA fragment encoding mIBP83-GFP was generated by overlapping extension PCR using pEX-A2-SBP-T and pGFP-cycle3 [10] (Melnik TN et al, PLoS ONE, 2012) with the following primers:

T7Prom 5’ CCCGCGAAATTAATACGACTCACTAT 3’;
mIBP83r 5’ GCCGCTCGCGCCCACGCCG 3’;
GFPf 5’ GCGAGCGGCACCGGCATGGCTAGC 3’;
T7Ter 5’ CTAGTTATTGCTCAGCGGTGGC 3’.

The amplified mIBP83-GFP fragment was inserted into pET28a vector (Novagen).

For all the constructs, the correct reading frames were confirmed by DNA sequencing.

### Isolation and purification of mIBP83 and mIBP83-GFP proteins

*E. coli* BL21 (DE3) strain was used for expression of ice-binding proteins. Transformation of competent E. coli cells was conducted using pET-22b plasmid encoding the target protein. Cells were cultivated in liquid LB medium of the following composition (per 1 L): trypton, 10 g; yeast extract, 5 g; NaCl, 10 g; NaOH (10 M), 600 µl. Kanamycin was added to the growth medium as a selection marker in 1:100 ratio, the gene of resistance to this antibiotic was encoded in the target vector plasmid. To stimulate the yield of the target protein, IPTG was added to the cultural medium IPTG to 100 mM final concentration. High-pressure homogenizer was used for cell disintegration. All the solutions were prepared using de-ionized and distilled water.

Protein isolation from *E. coli* cell lysate was conducted by sequential gel filtration, ion-exchange and hydrophobic chromatography. The purity of protein preparations was analyzed by SDS-PAAG in 12% gel. Concentration of ice-binding protein mIBP83 was determined by ultraviolet absorption at 280 nm, taking its extinction coefficient (0.683) into account. Concentration of GFP and mIBP83-GFP proteins was determined at 395nm wavelength corresponding to absorption band of the chromophore group of the green fluorescent protein.

Cycle3-GFP variant of green fluorescent protein was used in all the experiments [8, 9, 11]. It is named simply GFP in this article. Cycle3-GFP folds well at 37°C during expression in E.coli cells; that is why it is used in many studies as a “wild-type protein”. Besides, it is very important for the goals of the current work that this protein does not dimerize at low concentrations (up to 2 mg/ml) [8, 9, 11]. This was tested by sedimentation and analytical chromatography. mIBP83-GFP fusion protein was also analyzed by gel chromatography. The protein, as well as GFP, had an elution profile with one quite sharp peak, giving evidence that this protein does not dimerize.

### Instruments and materials used in the work

For fluorescence excitation in the experiments on interaction of GFP-mIBP83 protein with ice surface (Figure 4), ECX-F20.M transilluminator (Vilber Lourmat, France) was used.

The experiments on ice freezing-melting were carried out in an appliance device composed of PC-controlled thermostat (Julabo F-25, Germany). Temperature was measured by an electronic thermometer with 0.01°C specific accuracy (reproducibility) and 0.1°C absolute accuracy after adjustment around 0°C. The solutions were placed into 250 µl plastic tubes. The basic scheme of the instrument is shown in Figure 6.

**Figure 6.**
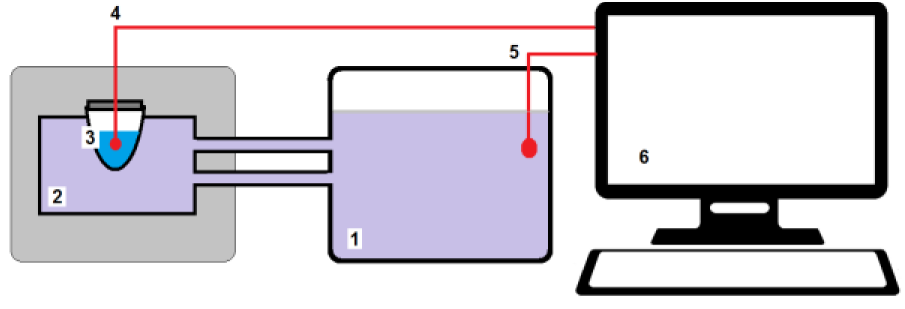
A device for melting and freezing the solutions. (1), thermostat; (2), thermally insulated cell, into which the sample (3) is placed. A sensor of the electronic thermometer (4) is immersed into the sample through tightly sealed cap of the tube. Sample temperature is also controlled by the thermometer (5). Personal computer (6) records the sample temperature and controls the linear change of thermostat temperature during heating or cooling.

## Acknowledgements

We are grateful to Alexei V. Finkelstein for fruitful discussions and Ekaterina N. Samatova for assistance. The work was supported by the Russian Foundation for Basic Research grant No. 19−04−00420.

## Author Contributions

B.S.M. conceived the idea for the study, experimental design, performed data analysis and wrote the paper; K.A.G - plasmids creating; B.S.M., J.D.O. performed the experiments.

## References

1 Graether SP, Kuiper MJ, Gagné SM, Walker VK, Jia Z, Sykes BD & Davies PL (2000) β-Helix structure and ice-binding properties of a hyperactive antifreeze protein from an insect. Nature 406, 325–328.

2 Leinala EK, Davies PL & Jia Z (2002) Crystal Structure of β-Helical Antifreeze Protein Points to a General Ice Binding Model. Structure 10, 619–627.

3 Kuiper MJ, Morton CJ, Abraham SE & Gray-Weale A (2015) The biological function of an insect antifreeze protein simulated by molecular dynamics. Elife 4.

4 Xu H, Perumal S, Zhao X, Du N, Liu X-Y, Jia Z & Lu JR (2008) Interfacial adsorption of antifreeze proteins: a neutron reflection study. Biophys J 94, 4405–13.

5 Tyshenko MG, Doucet D, Davies PL & Walker VK (1997) The antifreeze potential of the spruce budworm thermal hysteresis protein. Nat Biotechnol 15, 887–890.

6 Doucet D, Tyshenko MG, Kuiper MJ, Graether SP, Sykes BD, Daugulis AJ, Davies PL & Walker VK (2000) Structure-function relationships in spruce budworm antifreeze protein revealed by isoform diversity. Eur J Biochem 267, 6082–6088.

7 Garcia-Arribas O, Mateo R, Tomczak MM, Davies PL & Mateu MG (2006) Thermodynamic stability of a cold-adapted protein, type III antifreeze protein, and energetic contribution of salt bridges. Protein Sci 16, 227–238.

8 Glukhova KF, Marchenkov V V, Melnik TN & Melnik BS (2017) Isoforms of green fluorescent protein differ from each other in solvent molecules “trapped” inside this protein. J Biomol Struct Dyn 35, 1215–1225.

9 Fukuda H, Arai M & Kuwajima K (2000) Folding of green fluorescent protein and the cycle3 mutant. Biochemistry 39, 12025–12032.

10 Melnik TN, Povarnitsyna T V, Glukhov AS & Melnik BS (2012) Multi-state proteins: approach allowing experimental determination of the formation order of structure elements in the green fluorescent protein. PLoS One 7, e48604.

11 Melnik BS, Molochkov N V, Prokhorov DA, Uversky VN & Kutyshenko VP (2011) Molecular mechanisms of the anomalous thermal aggregation of green fluorescent protein. Biochim Biophys Acta 1814, 1930–1939.

